# A standardized head-fixation system for performing large-scale, *in vivo* physiological recordings in mice

**DOI:** 10.1101/2020.01.22.916007

**Authors:** PA Groblewski, D Sullivan, J Lecoq, SEJ de Vries, S Caldejon, Q L’Heureux, T Keenan, K Roll, C Slaughterback, A Williford, C Farrell

**Author notes:** Co-lead authors (PA Groblewski, D Sullivan).

## Abstract

**BACKGROUND:** The Allen Institute recently built a set of high-throughput experimental pipelines to collect comprehensive *in vivo* surveys of physiological activity in the visual cortex of awake, head-fixed mice. Developing these large-scale, industrial-like pipelines posed many scientific, operational, and engineering challenges.

**NEW METHOD:** Our strategies for creating a cross-platform reference space to which all pipeline datasets were mapped required development of 1) a robust headframe, 2) a reproducible clamping system, and 3) data-collection systems that are built, and maintained, around precise alignment with a reference artifact.

**RESULTS:** When paired with our pipeline clamping system, our headframe exceeded deflection and reproducibility requirements. By leveraging our headframe and clamping system we were able to create a cross-platform reference space to which multi-modal imaging datasets could be mapped.

**COMPARISON WITH EXISTING METHODS:** Together, the *Allen Brain Observatory* headframe, surgical tooling, clamping system, and system registration strategy create a unique system for collecting large amounts of standardized *in vivo* datasets over long periods of time. Moreover, the integrated approach to cross-platform registration allows for multi-modal datasets to be collected within a shared reference space.

**CONCLUSIONS:** Here we report the engineering strategies that we implemented when creating the *Allen Brain Observatory* physiology pipelines. All of the documentation related to headframe, surgical tooling, and clamp design has been made freely available and can be readily manufactured or procured. The engineering strategy, or components of the strategy, described in this report can be tailored and applied by external researchers to improve data standardization and stability.

## INTRODUCTION

One of the overarching goals of the Allen Institute for Brain Science is to deepen our understanding of the mammalian visual system, from the moment at which photons enter the eyes to the execution of complex visually guided behavior (Koch & Reid, 2012). To achieve this goal, we have constructed the *Allen Brain Observatories* to collect comprehensive maps of neural activity in awake behaving mice. These large-scale efforts will yield complementary optical physiology, electrophysiology, and behavioral datasets of unprecedented size and standardization—all of which will be made freely available to the scientific community for continued analysis and publication. Performing the experiments necessary to collect these datasets posed significant operational and engineering challenges. Here we describe the unique difficulties of, and our solutions for, collecting systematic physiological data from head-fixed mice at scale.

The *Allen Brain Observatory* pipelines were designed to progress mice through sequential data collection steps, each of which is performed by teams of technicians. A simplified overview of the experimental workflow of our first *Allen Brain Observatory* is illustrated in Figure 1 where each icon represents a distinct experimental step. Transgenic mice, such as those that express an activity-dependent fluorophore, GCaMP6 (Madisen et al., 2015), are bred and cared for in-house by our Animal Husbandry team before receiving surgery performed by our Surgical team. Following recovery, mice receive Intrinsic Signal Imaging by our Imaging team to obtain a functional map of the visual cortical areas. Mice are then transferred to our Behavior team for habituation prior to our Microscopy team that performs *in vivo* physiological recordings. Once *in vivo* experiments are completed, tissue is collected and all data is sent through our post-acquisition processing pipeline where data from multiple streams is extracted, transformed, assessed for integrity and quality, and, lastly, loaded into a shareable format. Data is made freely available to the public via our web portal. Here we focus on our first *Allen Brain Observatory* (shown in blue, Figure 1) which includes recordings of cortical activity from over 60,000 neurons collected from 6 visual areas, 4 layers, and 14 transgenic mouse lines from nearly 300 adult mice, in response to a systematic set of visual stimuli. To date, analysis of this initial dataset has been included in a number of publications by both Allen Institute researchers (e.g., Arkhipov et al., 2018; de Vries et al., 2019; Waters et al., 2019) as well as external researchers (e.g., Cai, Wu, & Ji, 2018; Esfahany, Siergiej, Zhao, & Park, 2018; Sweeney & Clopath, 2020). Future iterations of the Brain Observatory will include behavioral and physiology modality variants (shown in gray, Figure 1).

**Figure 1.**
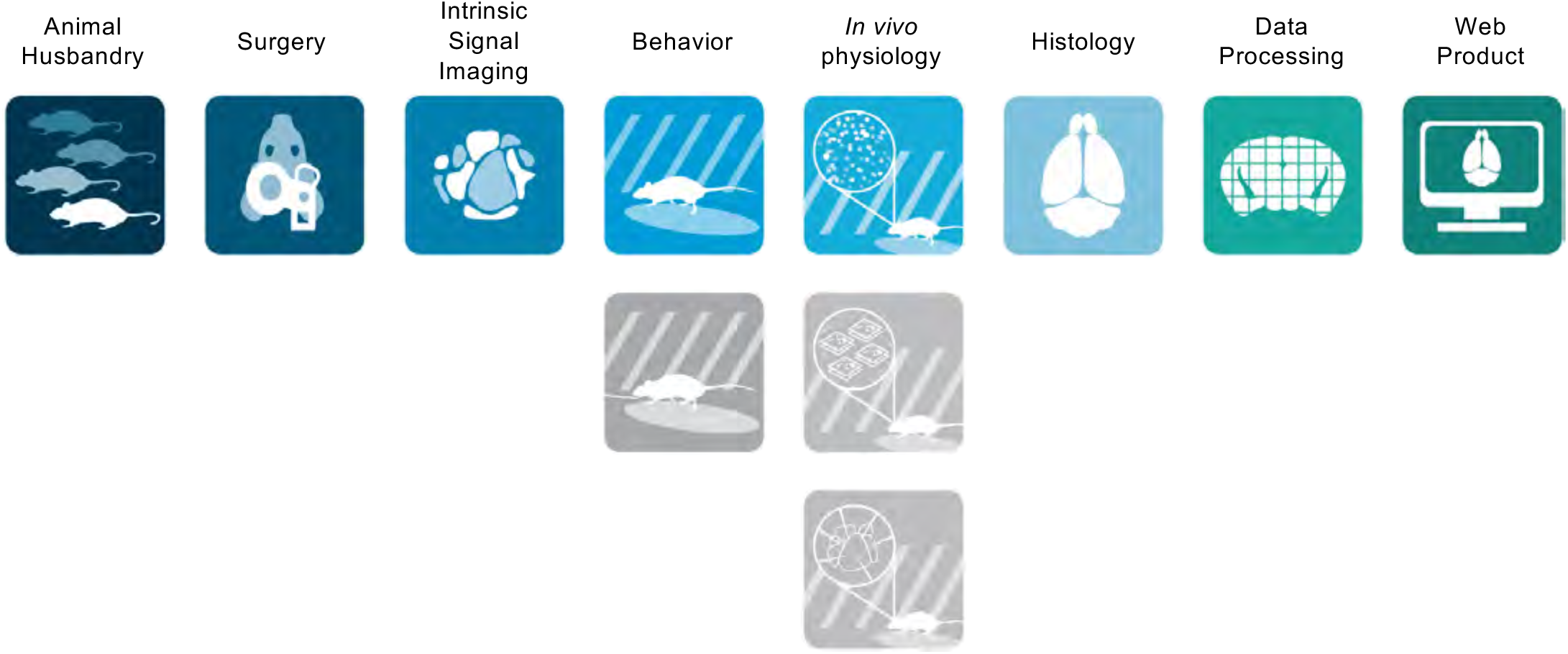
Allen Brain Observatory pipelines. Schematized view of the various Allen Brain Observatory pipelines. Each icon represents a distinct experimental step of the pipeline that is performed by a different team of technicians according to a series of standard operating procedures. Gray icons represent platforms that are part of subsequent Brain Observatories not detailed in this manuscript (including operant behavior training, multi-plane 2-photon calcium imaging, and multi-probe electrophysiology).

The operational and scientific requirements of the *Allen Brain Observatory* necessitated a comprehensive engineering strategy to ensure the stability and quality of datasets that were collected by teams of technicians over long-periods of time. Our strategy was to create a “cross-platform reference space” by defining and fully constraining datum structures on mating components—both in rotation and translation. This reference space is defined by the three mutually intersecting perpendicular datum planes created upon immobilization of the headplate within the clamp. Critically, this reference space allows for spatial information to be defined and translated both within and between platforms. Furthermore, it allows consistent placement of all experimental components and instrumentation relative to the reference space and therefore to the mouse.

The fundamentals of a cross-platform reference space are based on the definition of a shared Cartesian coordinate space that is referenced to known biological features; in this case we relied on common mouse skull fiducials including lambda, bregma, and the interaural line. There are three components that are necessary for creating a shared coordinate space for data collected from hundreds of head-fixed mice using many instruments: 1) a robust headframe, 2) a reproducible clamping system, and 3) data-collection systems that are built, and maintained, around precise alignment with a reference artifact.

Here we describe in detail the engineering strategies and tools we implemented to build and maintain the *Allen Brain Observatory* pipelines. Our strategy for melding standardization with scale was to create an experiment-wide coordinate system, or cross-platform reference space. The cross-platform reference space required designing and validating a suite of equipment and tools, which we describe below and are being made freely available as an open resource to the scientific community.

## RESULTS

The first engineering challenge that we faced in building a cross-reference space for the *Allen Brain Observatory* pipelines was to design a robust headframe. Together with our scientific and operational teams we defined the following headframe requirements:

1. <u>Registrability and rigidity:</u> the headframe needed be rigid (no more than 4 μm of deflection) and registerable across various instruments, allowing for a single set of cells to be recorded by multiple instruments over many experimental sessions.
2. <u>Basic geometrical and experimental constraints:</u> the headframe and well needed to be compatible with our various pipeline instruments and allow for physiological recording access to all visual cortical regions.
3. <u>Animal health:</u> the headframe needed to be lightweight (no more than 10% of bodyweight), made of biocompatible material, and not impair normal in-cage mouse behavior (including locomotion, feeding and drinking).
4. <u>Ease of handling:</u> head-fixation needed to be easy and quick, thereby reducing unnecessary stress on the animal and experimenter.
5. <u>Adaptability</u>: the headframe and clamp design needed to be adaptable to other types of physiological recordings from various brain regions.

We performed an assessment of the current (as of 2014) state of the art including designs used by various laboratories (e.g. Andermann et al., 2010; Guo et al., 2014) as well as commercially available options (e.g., https://www.neurotar.com/). We determined that none of these <u>individual</u> headframe designs satisfied <u>all</u> of our requirements. For instance, most existing headframes required two holding clamps—a design that would force the system to be statically indeterminate, thereby preventing knowledge of exact headframe location (and thus any cells of interest) as well as placing physical strains on the headframe. Furthermore, most headframe clamp interfaces lacked constrainable datum structures necessary to remove rotational and translational degrees of freedom (required for reproducible placement of the headframe on the skull, as well as registration across instruments/platforms). Upon assessment of existing headframes, we decided to create a novel design, which allowed us to incorporate all the necessary features from the ground up. While iterating on the headframe design we co-developed a surgical procedure and set of custom surgical tooling that would precisely place the headframe relative to the mouse skull. (*See Headframe and Headframe Surgical Tooling)*

The second requirement of a cross-platform reference space is a robust clamping system and for this we relied on a clamping interface that we had incorporated into the headframe shank. The headframe’s built-in registerable faces/features allowed it to be reproducibly placed and secured into the clamps of the various instruments. *(See Clamping System & System Alignment)*

The final requirement of the cross-platform reference space is that the data collection systems are designed, built and maintained around precise alignment with a single reference artifact; we used a reticle. An important challenge that we faced was that the pipeline platforms are diverse and possess unique sets of constraints related to both the mode of data acquisition (e.g. ISI camera vs 2-photon microscope objective) and physical attributes (including size, shape, and orientation). To standardize mouse placement across these diverse platforms we created hardware solutions that incorporated a common mouse stage that placed mice at a fixed geometry with respect to the visual stimulus monitor (Supplemental Figures 1, 4, and 5). Importantly, these features are set and validated independently of biological variation and permit long-term monitoring of system alignment. *(See Systems & Applications and Cross-Platform Registration)*

### Headframe

#### DESIGN

The headplate design and dimensions are shown in the plan and side views of Figure 2a, and isometric views of Figure 2b. The headplate comprises two main parts: a shank (the feature that is loaded into a clamp) and the mouse-interface that, in this case, encircles the visual area of the cortex in the left hemisphere (see below for stereotaxic coordinates).

**Figure 2.**
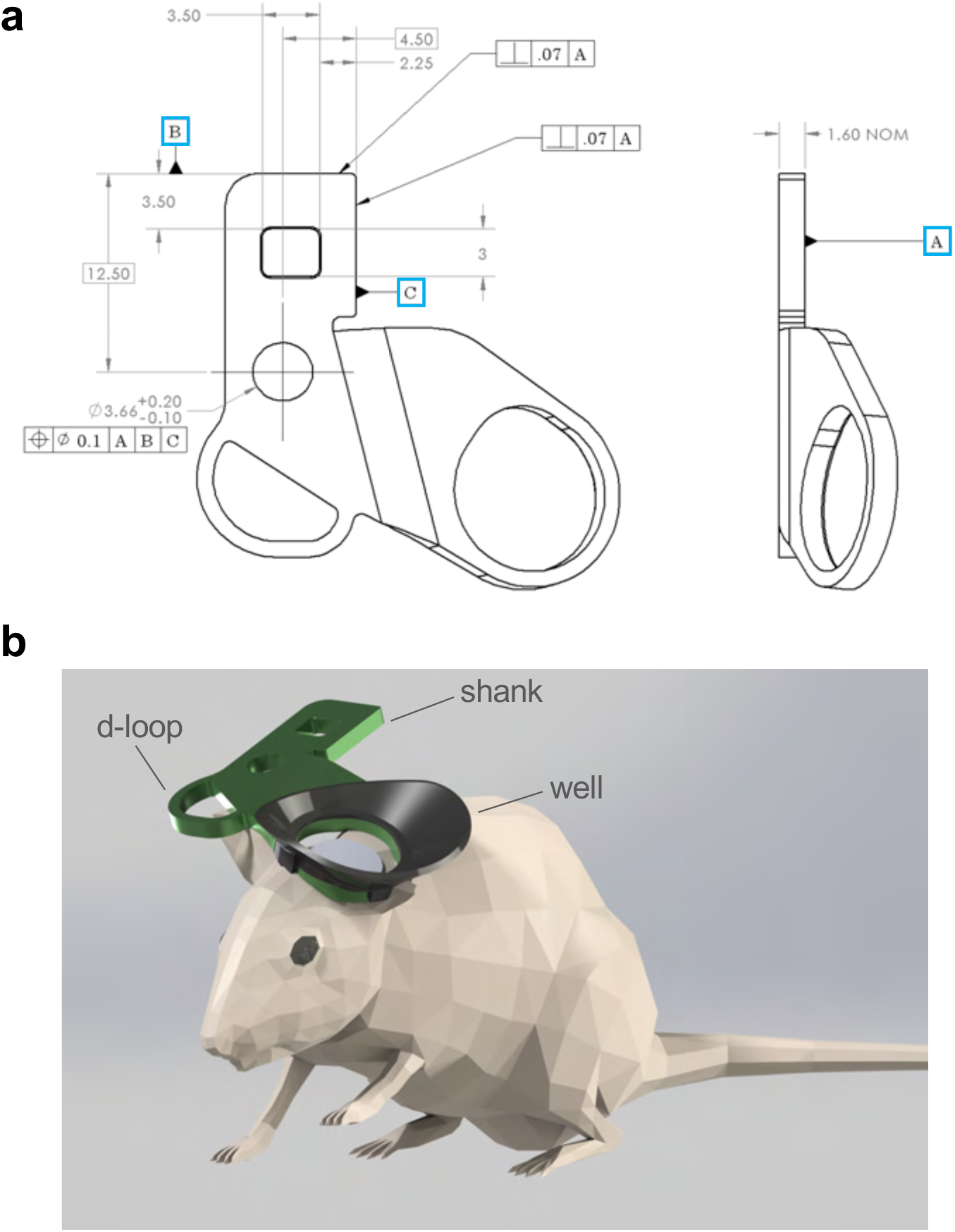
Brain Observatory headframe. a) Plan- and side-view of the Brain Observatory headplate including dimensions. The three datum reference planes are shown as A, B, and C. Headplates are manufactured from 1.6 mm (.063 inch) thick titanium 6Al-4V using common manufacturing methods. b) Isometric view of headframe affixed to a mouse. Headplate shank projects posteriorly so as to not obstruct the mouse’s field of view or impede on the mouse’s ability to locomote.

The shank design includes three perpendicular, manufactured surfaces that define a datum reference frame (shown as A, B, and C in Figure 2a). The reference frame on the headplate mates to the corresponding reference frame on the clamp of each instrument and constrains the three translational (x, y, z) and three rotational (pitch, roll, yaw) degrees of freedom. The common reference frame provides a physical origin point to which all visual stimuli and instrumentation are placed in three-dimensional space. Incorporating the physical origin point in the design of each of our instruments ensures that the mouse experiences the same stimulus across every platform (and every instrument within each platform) of the pipeline.

The design of the mouse interface of the headplate was driven by 1) the area of interest for physiological recordings, 2) the microscope or other instrument interface, and 3) the geometry of the mouse skull. The headframe developed for the *Allen Brain Observatory* consists of a 10 mm circular opening centered over the putative location of the primary visual cortex (M/L= -2.8 mm, A/P = 1.3 mm, with respect to lambda). The 10 mm ring features a slight teardrop shape to accommodate skull variation between animals and provides clearance for a more reproducible, unobstructed skull contact point across mice. The headplate is mated to a water-retention well that was designed to interface with a 16X Nikon CFI LWD Plan Fluorite Objective.

A final requirement of the headplate was that it should provide a finger-hold to help ease the clamping procedure for head-fixation. The ‘d-loop’, visible in Figures 2a and b, fulfills this need without adding significant weight, requiring additional handling tools, or obstructing the mouse’s view of the stimulus monitor.

#### ANIMAL HEALTH REQUIREMENTS

The *Allen Brain Observatory* headplate is manufactured from 6Al-4V titanium using common manufacturing methods *(see Materials & Methods)*. Titanium’s biocompatibility is well understood and, as such, it is often selected for implants in mice and humans (Sidambe, 2014). Given its desirable stiffness-to-weight ratio it is an especially good fit for this application and is routinely used for similar *in vivo* neuroscience applications (e.g., Guo et al., 2014; Hefendehl et al., 2012). The finished headplate mass is ∼1.9 g (approximately 10% of body weight of a minimum weight mouse at the time of implantation). We monitored in-cage behavior of mice following surgery and observed that mice can eat, drink, and locomote normally with the headframe and well attached to the skull. Given our observations that the headframe had no observable impact on normal animal health and behavior we do not specifically track this in an experimental setting. However, as part of our initial *Allen Brain Observatory* pipeline (de Vries et al., 2019), we reported, approximately 14% of mice were excluded from the final dataset due to a health issue that may have been surgery-related; that is, not directly caused by previously observed phenotypic issues (e.g., dermatitis, epilepsy) specific to the transgenic mouse lines (http://observatory.brain-map.org/visualcoding/transgenic). It is important to note, however, that these exclusions were largely due to the invasive nature of the cranial window portion of the surgery, and not the headframe itself.

#### EXPERIMENTAL REQUIREMENTS

The experimental paradigm requires that, to the greatest extent possible, the mouse’s view of the stimulus screen is unobstructed. The visual field of the right eye extends approximately to 110 degrees vertical, 140 degrees horizontal (Wagor et al., 1980) while the stimulus monitor extends to +/-47 degrees in the vertical and +/-59.3 degrees in the horizontal from the gaze axis of the mouse. To keep the visual field clear, the shank of the headplate was kept to the rear of the skull and was also raised above the body of the mouse to keep it clear of the neck and back (that is, it is parallel to and elevated from the x-y plane of the mouse coordinate system) (Figure 2b).

As one of the experimental requirements for our headpost design, our 2-photon (2P) microscopy team determined *a priori* that the imaging region of interest must exhibit no more than 4 μm of total displacement along the optical axis during an experiment. Figure 3a shows the results of a computer simulation to assess how the headframe and clamp performed under a downward, 0.5 N load distributed across the entire ring of the headplate. We deemed a 0.5 N force to be a conservative estimate when compared to an average running mouse with mass of approximately 20 grams (0.196 Newtons) without the ability to exert significant force on a low stiffness, foam-covered running disk. The simulations showed that the 0.5 N load resulted in a displacement of approximately 1.5-2 μm at the center of the ring, with a maximum deflection of ∼3 μm at the outer edge of the ring. Furthermore, the simulations revealed that the deflection was limited to the headframe itself, and not the shank or clamp. Figure 3b shows the results of our bench tests during which we measured the actual deflection (measured at ring center and rim using a laser displacement sensor) caused by 20 g, 50 g, and 100 g weights hung under the outer rim of the headplate ring. Our tests confirmed that the titanium headpost, coupled with our pipeline clamping mechanism, demonstrated an average of 3.2 μm of displacement at the center of the ring under a static 0.49 N (50 g) load applied to the far end of the headplate. The displacement observed in the bench test is likely limited to the headframe ring, given the similarity of these results to the simulations. A final set of tests were performed to assess the deflection of the headframe and cranial window surface in actual mice. Specifically, following recovery from the Headframe & Cranial Window surgery (see *Materials & Methods*) two mice were head-fixed in a pipeline 2P microscope instrument and deflection measurements were obtained while the mice were actively locomoting (i.e., during bouts of running and stopping). The results of these tests are shown in Figure 3c, overlaid with a baseline measurement of noise. As expected, there was minor displacement of the headframe when the mice were locomoting and this displacement increased slightly when measured at the surface of the cranial window (as indicated in the example traces and histograms, as well as reflected by the increase in signal variance). Importantly, however, maximum displacement (≤ 2.2 μm), and even total range of displacement during the entire recording period (≤ 3.3 μm), was well within our experimental tolerances.

**Figure 3.**
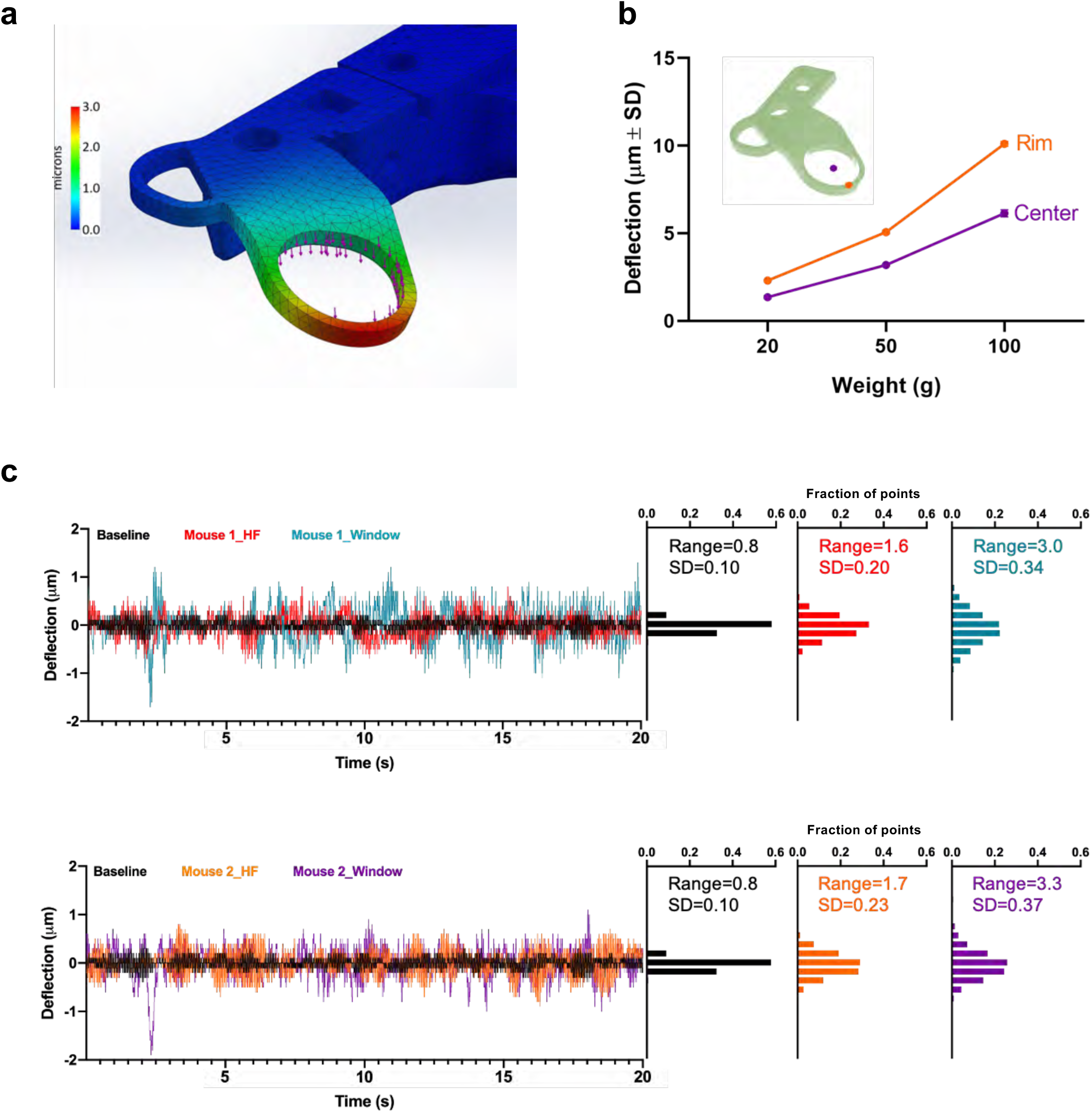
Headframe stiffness testing. a) Deflection simulations of the headplate and clamping mechanism suggested that a static .5N load applied to the headplate ring (shown as purple arrows) resulted in ∼1.5-2 μm deflection at the center of the ring and a maximum of ∼3μm deflection at the outer edge of the ring. b) Results of benchtop deflection tests indicated that a static load of 50g (∼0.5N) applied to the rim of the headplate resulted in an average of 3.2 μm of displacement at the center of the headplate ring. c) Deflection was next measured on the headframe (HF) and cranial window surface (Win) of two running mice that were head-fixed in an optical physiology instrument (Supplemental Figure 4a). Twenty-second samples of deflection data obtained from each point are shown, overlaid with the baseline noise measured on a headframe only (Baseline) clamped into the same system. As expected, deflection of the headframe during running increased over baseline and was greatest on the cranial window, as indicated by the increases in variance of the recorded signal (further shown as a broadening of the frequency distributions). However, the maximum deflection distances and ranges were well within our pre-determined tolerances.

**Figure 4.**
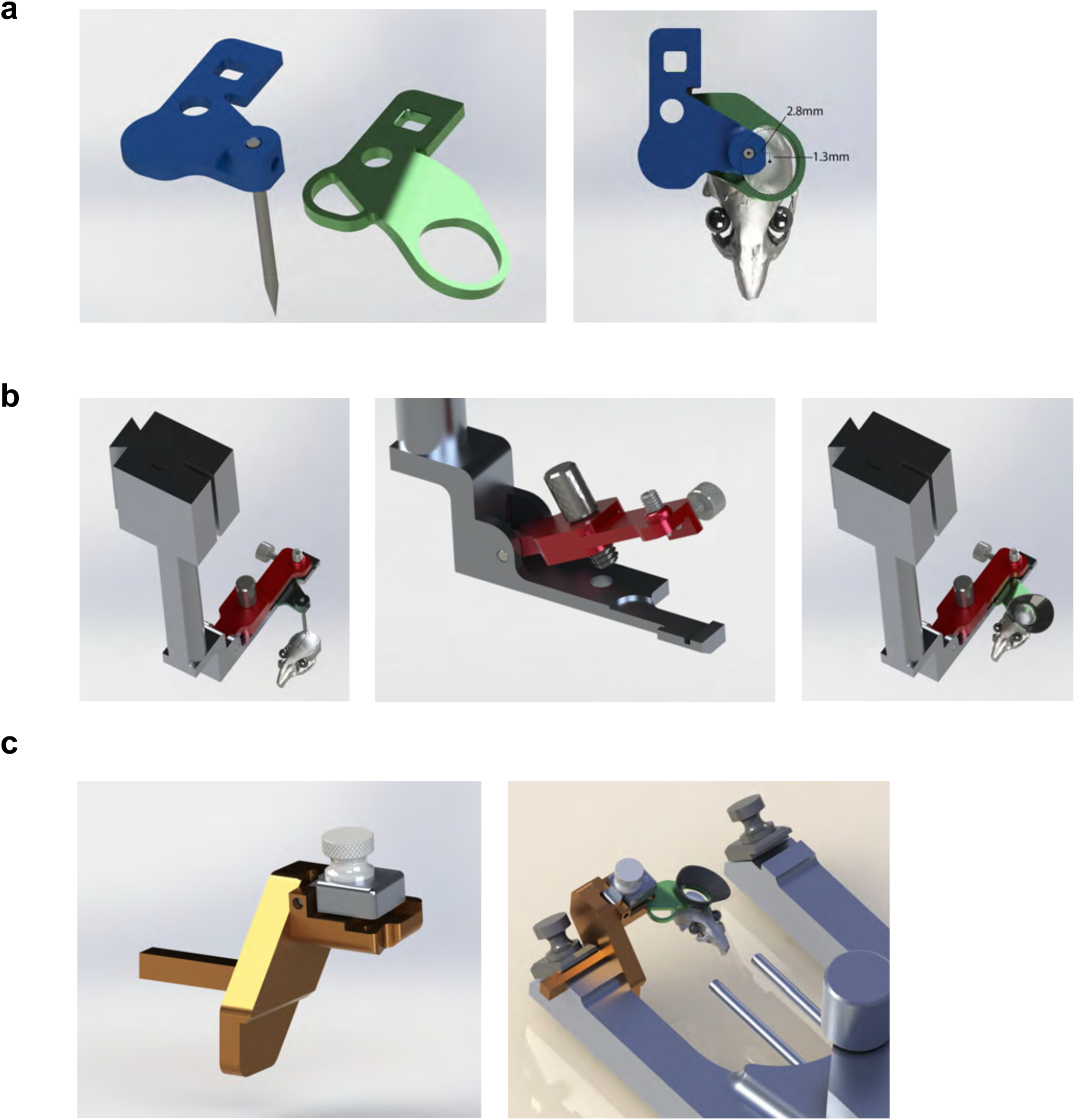
Headframe surgical tooling. a) LEFT: A “stylus” (blue), equipped with a shank identical to that of the headplate (green), is used to locate the mouse skull fiducial, lambda. RIGHT: This stylus places the headplate so that the center of the 10 mm headplate ring is located over the putative location of mouse visual cortex (M/L = -2.8mm, A/P = 1.3mm, with respect to lambda). b) Custom headframe placement tool (“headframe clamp”) is compatible with the KOPF 1900 dovetail interface located on the Z-axis arm and places a clamshell-style clamp (CENTER) parallel to the levelled mouse skull. Once lambda is located with the stylus (LEFT), it is replaced by the headframe (RIGHT) and lowered down along the Z axis to the skull for cementing. c) The cranial window portion of the surgical procedure is facilitated by a custom “levelling clamp” (LEFT) that interfaces with the KOPF 1900 earbar clamp (RIGHT) and pitches the mouse forward 6 degrees. Once the entire earbar apparatus is rotated to 23 degrees, the cranial window plane is positioned perpendicular to gravity.

### Headframe Surgical Tooling

The *Allen Brain Observatory* cranial window surgery has previously been described in detail (de Vries et al., 2019) as well as in the *Materials & Methods*. Additional information can be found at https://help.brain-map.org/display/observatory/Documentation.

#### HEADFRAME PLACEMENT

To standardize the placement of the headframe onto the mouse skull we designed a custom set of tools that remove all angular degrees of freedom, as well as X and Y variability, from the headframe installation process. These tools include a “headframe clamp” and “stylus”. The “headframe clamp” tool interfaces with KOPF Model #1900 dovetail mounts and suspends a clamshell-style clamp above the mouse. Once the mouse’s skull is levelled with respect to pitch (bregma-lambda level), roll, and yaw, the “headframe clamp” tool is secured in the KOPF arm and aligned so that the clamp is positioned at a known offset from lambda (the coordinate system origin). This is achieved with the custom “stylus”, shown in Figure 4a. The “stylus” tool shares the same shank as the headplate and so utilizes the same datum registration surfaces. It is installed in the “headframe clamp” tool (Figure 4b) and the surgeon adjusts the X and Y axes of the stereotaxic instrument to locate the tip of the “stylus” at lambda. Once lambda is located, the “stylus” is removed from the clamp and a headframe is installed and lowered until the anterior portion of the headplate contacts the skull. The headframe is cemented to the skull and once dry, the clamshell is opened, releasing the headframe. This tool reproducibly places the headframe such that the center of the well is 2.8 mm lateral and 1.3 mm anterior to lambda (Figure 4a).

#### CRANIOTOMY & CRANIAL WINDOW

To facilitate repeatable location of the craniotomy and cranial window we designed a custom clamp pictured in Figure 4c. The “levelling clamp” was designed to fit into the KOPF Model #1900 earbar holder upon removing the right earbar and is compatible with the anesthesia nose cone (although the mouse must be removed from the bite bar clamp). The “levelling clamp” has a built-in forward pitch of 6°, and along with rotation of the entire earbar apparatus to a roll angle of 23° (using a custom-adapted angle finder not pictured) it holds the craniotomy plane perpendicular to gravity. Once the headframed animal is clamped and rotated, a circular piece of skull (5 mm in diameter) is removed with a dental drill, and a durotomy is performed. The “levelling clamp” facilitates drilling of the craniotomy (and subsequent durotomy) by 1) allowing the surface of the skull (and subsequently the brain) to be more clearly viewed through the stereo microscope and 2) keeping artificial cerebrospinal fluid used during the procedure contained within the headplate ring. Following the craniotomy and durotomy, a 0.45 mm thick custom borosilicate glass coverslip (stacked appearance with a 5 mm diameter “core” and 7 mm diameter “flange”) is cemented in place. The “leveling clamp” facilitates consistent placement and cementing of the cranial window at an angle that is approximately parallel to the headplate ring, and thus normal to the imaging axis of our pipeline data-collection instruments.

### Clamping System & System Alignment

#### CLAMP DESIGN

To accurately place the animal relative to stimulus monitor and instrumentation (e.g., the microscope objective) we developed a custom clamping mechanism shown in Figure 5. This clamping mechanism simultaneously meets multiple requirements including 1) reproducibility, 2) high stiffness, 3) quick installation and removal with common tools, and 4) compatibility with all *Allen Brain Observatory* platforms. An additional design requirement was that the clamping mechanism be manufactured with commonly available screws, materials, and processes. The clamp was designed to position the headplate shank into the common datum surface with two, screw-driven mechanisms pushing perpendicular to each other and at 45 degrees with respect to the planes they are pushing against (see Figure 5a). The headplate is inserted, and positively located into the corner of the clamp and it can be installed or released in under 10 seconds (Figure 5b). An optional third screw (Figure 5b) is utilized for applications demanding the utmost stability of the animal (e.g., 2P microscopy). The datum surfaces of the headplate and clamp are broad to prevent wear, while the force application points can accommodate manufacturing variation and wear without sacrificing clamping accuracy. It is important to note that accuracy is highly dependent on clean reference surfaces; buildup of dirt, debris and animal dander will impact clamping accuracy and, therefore, headplate cleanliness must be maintained throughout the duration of experimentation.

**Figure 5.**
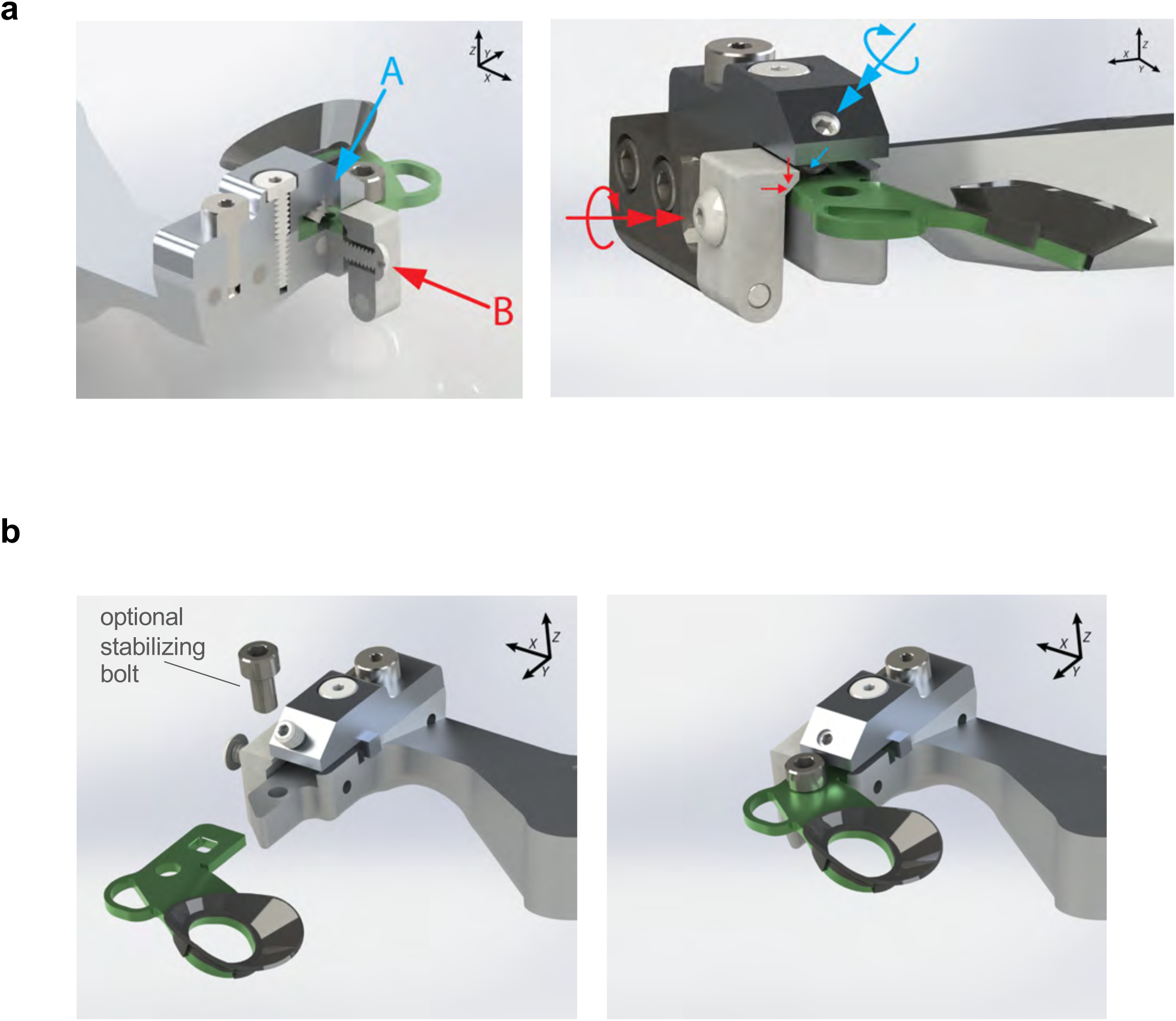
Headframe clamping system. a) Sectioned isometric view of the headframe clamping system exposes the two clamping bolts (A and B) that ensure proper seating of the headframe to the clamp. Two axial screw forces are applied to the headframe during installation. Red arrows show that axial screw force on the side bolt in the X direction resolves to both Z and X forces applied to the outer edge of the headframe shank and applies clamping pressure to datums A and C (Figure 2a). Blue arrows indicate that the axial screw force applied at 45 degrees in Z and Y resolves in the same forces and clamping pressures applied to the headframe in datums A and B. b) The headframe is inserted into the clamp along the Y axis and once clamped, can be further secured using the optional socket head cap screw situated anterior to the front face of the clamp.

#### SYSTEM ALIGNMENT

Despite being built around a common mouse-to-screen geometry (depicted in Supplemental Figure 1a), each of the *Allen Brain Observatory* data-collection systems possessed different rotational, translational, and scaling attributes of image acquisition. Furthermore, each system included multiple components that were each defined in unique coordinate spaces. An example of this is shown in Supplemental Figure 2, which depicts the various coordinate spaces that exist in the 2P imaging systems. Because of this, any X, Y, Z shifts in one of these spaces (e.g., the headframe) necessarily results in a different X, Y, Z translation in the others (e.g., the microscope). Therefore, in order to align all of our pipeline systems we employed a registration artifact in the form of a reticle that incorporated the geometry and clamping interface of the experimental headframe (Figure 6a). The reticle was permanently mounted in the headframe at approximately the imaging depth of interest and normal to the optical axis. Alignment of the system (either initially or during maintenance) involved initial mechanical alignment of the objective axis followed by fine alignment along the reticle imaging plane by obtaining instrument-specific reticle images at high magnification. Focus and rotation were defined by five points on the reticle, which enabled establishment of the imaging plane and a reticle center home position in system-specific stage coordinates. Alignment of the reticle home and defined imaging plane allows the user to navigate within the cortical layers that are parallel to the cranial window using only planar input (even though these 2-dimensional movements required the stage to move along all 3 axes). Exact positioning of all reticle points was monitored over time to ensure consistent system registration (see *Systems & Applications*).

**Figure 6.**
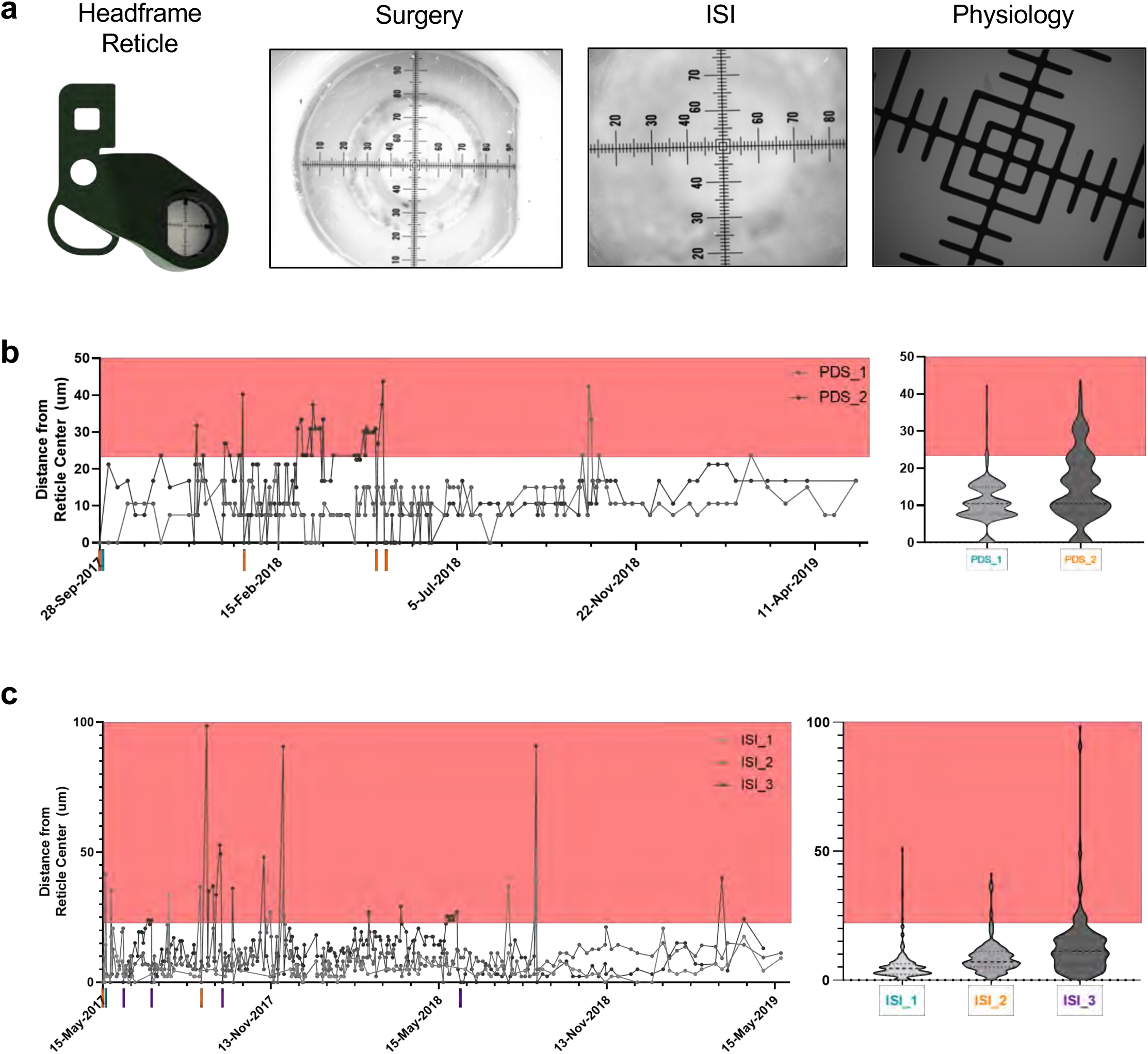
Pipeline system alignment and registration. a) Systems were aligned to a set of headframe reticles. Example images from photodocumentation, intrinsic-signal imaging, and 2P microscopy instruments that were used for initial alignment and longitudinal monitoring. The images highlight the unique rotational, translational, and scaling attributes of each of the platforms. b) Photodocumentation system registration monitoring data is shown for 2 instruments (PDS_1 & 2) over an 18-month period. Repeated detections of >22.5 μm deviation from the registered reticle location triggered system re-registration (shown as colored hash marks). Median clamping variability over this time period was 10.61 μm. c) Intrinsic-signal imaging system registration monitoring data is shown for 3 instruments (ISI_1-3) over a 24-month period. Detections of >22.5 μm of deviation from the registered reticle location triggered system re-registration (shown as colored hash marks). Median variability ranged from 4.5 to 11.25 μm depending on ISI instrument.

It is worth noting that alignment of a small number of instruments can be performed with a single reticle. However, in our case we implemented a second layer of abstraction wherein instruments are registered to a platform-specific, secondary reticle. Each of the secondary reticles is registered to a single primary reticle. Thus, images obtained on an instrument were translated to a common primary coordinate space using that instrument’s platform’s secondary-to-primary set of translation values. A two-layer reticle system allowed us to independently maintain alignment of 40+ instruments across 4 platforms without having to rely on a single reticle.

### Systems & Applications

#### SURGICAL PHOTO-DOCUMENTATION

Post-surgical brain health and window clarity were documented using a custom surgical photo-documentation system (in addition to normal animal health checks at one, two, and seven days following surgery). The photo-documentation apparatus (Supplemental Figure 3a) was custom designed to provide a registered image of the cranial window using the standard pipeline geometry. Because mice were imaged at the end of surgery, they were still lightly anesthetized and, as such, there was no need to include the third screw in the clamping mechanism. Additionally, because there was no visual stimulation for this data-collection step, the system did not include a stimulus screen.

Each of two photo-documentation systems were initially registered, and subsequently monitored weekly, by analyzing images of a secondary reticle. A sample of 18 months of longitudinal registration monitoring data is shown in Figure 6b. Repeated detections of >22.5 μm deviation from the registered reticle location triggered system re-registration (shown as colored hash marks). The median of monitoring data for each system was calculated and indicated a clamping variability of 10.61 μm for these systems. Thus, the photo-documentation clamping system exhibited stable clamp registration over long periods of time and required only periodic re-alignment.

#### INTRINSIC SIGNAL IMAGING

The *Allen Brain Observatory* pipelines utilize intrinsic signal imaging (ISI) with every mouse for targeting physiology recordings. Briefly, ISI measures the hemodynamic response of the cortex to visual stimulation across the entire field of view in mice that are lightly anesthetized. This retinotopic map effectively represents the spatial relationship of the visual field to locations within each cortical area. Retinotopic mapping is used to delineate functionally defined visual area boundaries and enable targeting of the *in vivo* physiology to retinotopically defined locations in primary and secondary visual areas (Garrett et al., 2014).

The ISI instruments had a different mouse-to-screen geometry (compared to the other pipeline platforms) and comprised an Andor Zyla 5.5 sCMOS camera and a ring illumination system of independently controlled green and red LEDs (Supplemental Figure 3b). The camera was fixed normal to the nominal window pitch and roll (6° and 23°, respectively). In addition to the camera and stimulus screen, the ISI system was equipped with an anesthesia machine (SomnoSuite, Torrington, CT) that was used to maintain a light plane of anesthesia during the ISI session. Because mice were lightly anesthetized there was no need to include the third screw in the clamping mechanism. As with the surgical photo-documentation system, each of three ISI systems were monitored for registration using a secondary reticle. A sample of 24 months of longitudinal monitoring is shown in Figure 6c. Colored hash marks on the x-axis indicate when the different systems required re-registration (due to reticle measurements drifting more than 22.5 μm from the original location). The median for each system indicated ISI clamping variability ranging from 4.5 to 11.25 μm. Therefore, as with the PDS system, the ISI system exhibited stable clamp registration over extended periods of time and required only periodic re-alignment.

#### BEHAVIOR

To support multiple versions of the *Allen Brain Observatory* we designed and built a large-scale behavior training facility that could simultaneously accommodate the behavior-training requirements of multiple pipelines. Each of these mouse behavior training enclosures (24 in total) were identically built to maintain the standard pipeline mouse-screen geometry.

The engineering requirements for the behavior platform included 1) a compact, modular design that allowed for training of ∼100 mice per day, 2) the ability to perform several different behavior tasks in different enclosures concurrently, 3) easy and reproducible clamping, and 4) pipeline mouse-screen geometry. Supplemental Figure 4a shows a front view of a behavior enclosure equipped with stimulus screen, sound-attenuating foam, ventilation fan, and a fixed-location camera (Allied Vision, Mako G-032B) to continuously monitor mice while in the enclosure. Mice are head-fixed on a removable behavior stage equipped with a running disc (Supplemental Figure 4b) and then placed onto a kinematic mount in the behavior enclosure, thereby ensuring quick but reproducible placement of the mouse with respect to the screen.

#### IN VIVO, 2-PHOTON CALCIUM IMAGING

The first iteration of the *Allen Brain Observatory* pipeline consisted of *in vivo* 2P calcium imaging in awake mice over multiple sessions/days. Our pipeline data collection systems were built around two off-the-shelf microscope models, Scientifica Vivoscope or Nikon A1R MP+, that we modified to accommodate our scientific and engineering requirements for pipeline data collection. In addition to incorporating our behavior stage (with running disc) and stimulus screen (ASUS PA248Q), each system was equipped with eye-tracking and full-body cameras (Allied Vision, Mako G-032B), each with their own LED illumination source. Both the Nikon and Scientifica systems are shown in Supplemental Figure 5.

The engineering requirements for the 2P calcium imaging platform were the most stringent and included 1) the ability to navigate to the same 400 × 400 μm field of view (and thus the same neurons) over multiple sessions/days, 2) a stable, rigid headframe and clamping system that allowed for no more than 4 μm of flex along the optical axis, and 3) pipeline mouse-screen geometry. Clamping performance was tested in an experimental context and the results of these tests are reported below.

### Cross-Platform Registration

As mentioned previously, to accommodate inter-instrument variability, we employed a reticle registration procedure that ensured initial system alignment as well as maintenance of that alignment over time. Precise reticle alignment had the added benefit that image capture from a diverse set of platforms could be translated to, and compared within, a shared coordinate space through a series of rotational, axial, and scaling factors. Specifically, each instrument had an established set of translation values to a platform-specific secondary reticle (obtained during system alignment, monitored, and updated if necessary), and each secondary reticle had a known set of different translation values to a common, primary reticle. As such, images obtained with any of our pipeline instruments could be translated to a common image space, resulting in “cross-platform registration” (Figure 7a).

**Figure 7.**
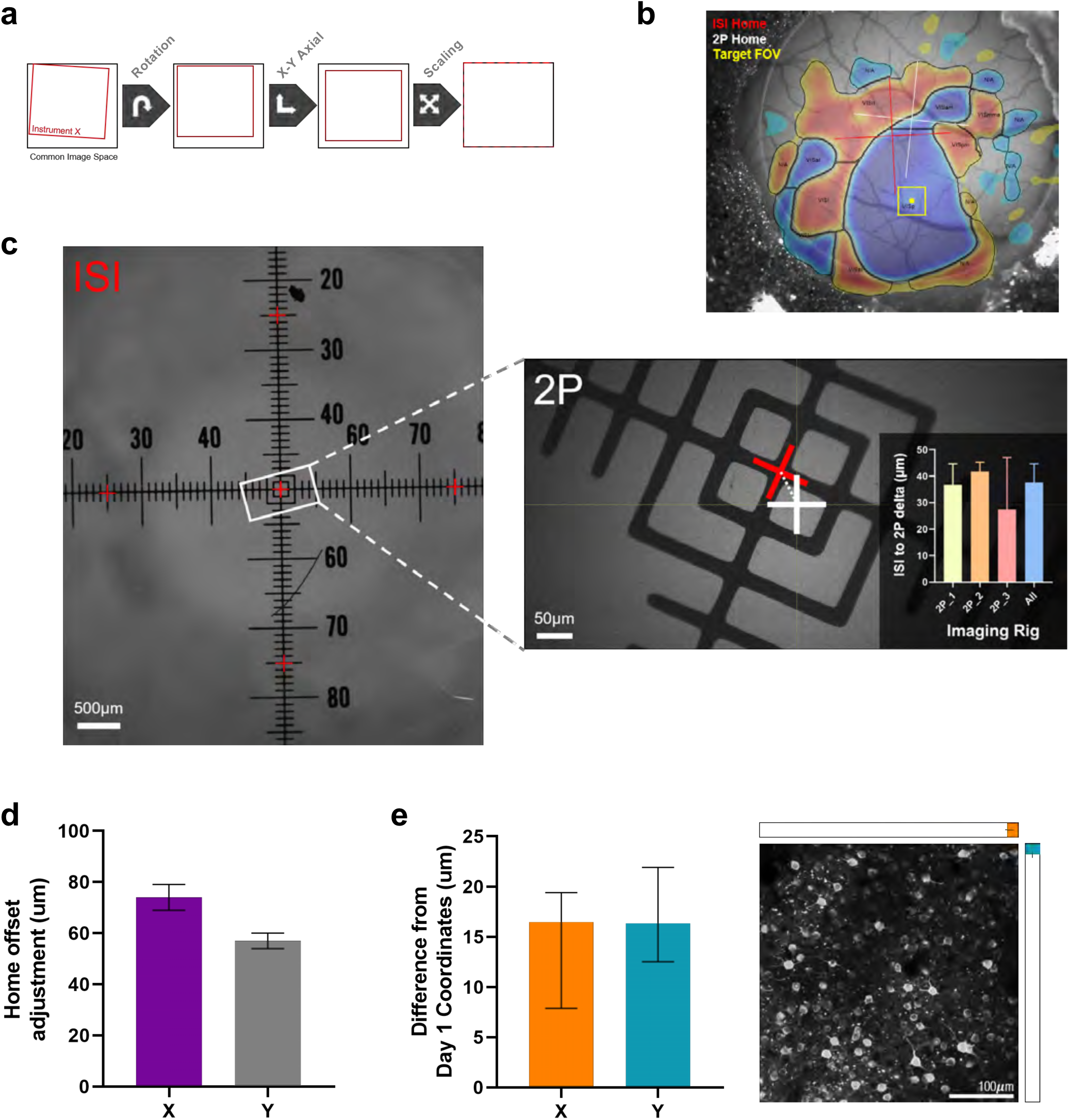
Cross-platform registration. a) Cross-platform registration relied on transforming data collected with an individual pipeline instrument to a common reference space. These translations included rotation, X-Y axial, and scaling factors that were specifically calculated for each instrument during the reticle alignment process. b) The *Allen Brain Observatory* leveraged cross-platform registration by using a set of ISI-defined coordinates to target 2P data collection of a retinotopically-defined region of cortex. c) Reliability of ISI-2P registration is shown using a non-biological sample. Trial-to-trial variability of navigating to a set of 5 ISI-translated coordinates (red crosses) on multiple 2P microscopes. Across the three pipeline 2P rigs targeting was within 10-50 μm (median of “All”= 37.6 μm) of the desired location. d) If necessary, before beginning an experiment operators were able to adjust the 2P home offset to adjust for biological motion and/or targeting inaccuracy. Median X and Y adjustments from a sample of >1700 experiments were 74 and 57 μm, respectively. e) LEFT: Experimental data depicting the X and Y adjustments (median and 95% CI) that operators made to match a 2P field of view (FOV) across sessions. Operators made ∼16um (median) of adjustment in both directions in order to match the FOV from the first session. RIGHT: An example 2P FOV (400×400 μm), with the median X (orange) and Y (teal) adjustments shown to scale.

The *Allen Brain Observatory* pipelines utilize ISI maps obtained from each individual mouse to perform physiological recordings in precise, retinotopic locations within the visual cortex. Because the instruments of the ISI and 2P microscopy platforms were all registered to a common reference space, we were able to identify the coordinates of a retinotopically-defined region (referenced to the ISI home location), translate the coordinates to the 2P reference space, and then drive to the target recording location from the 2P home location (Figure 7b). Accuracy of the ISI-2P translation in a non-biological sample is depicted in Figure 7c, which shows the trial-to-trial variability of navigating to a set of five ISI-translated coordinates (red crosses) on various 2P microscopes. Across three pipeline 2P instruments, targeting was 10-50 μm (median = 37.6 μm, sd = 9.93) off from the desired location, where factors contributing to this variability included clamping and coordinate translation.

In an experimental setting, however, there are additional biological factors that can impact targeting accuracy, including brain motion. Prior to each 2P recording session, operators were able to make an adjustment of the 2P home location so that it matched the ISI home location; this was performed in epifluorescence mode with 800 μm field of view (FOV). This “home offset” was then applied to the translated X and Y coordinates of the target recording location chosen from the ISI map. In a sample of our pipeline experiments (1712 sessions), the “home offset” adjustment values ranged from 0 to 905 μm, with median values of 74 μm and 57 μm in X and Y, respectively (Figure 7d). As previously mentioned, this adjustment accounted for variability caused by brain motion and was often necessary in order to adjust for any tissue movement that had occurred since the ISI, or previous 2P, session.

After navigating to the target recording site using the translated ISI coordinates, the operator was able to make final adjustments to the 400 × 400 μm, 2P FOV to ensure optimal cell matching with previous recording sessions. Figure 7e shows the day-to-day FOV targeting results from a sample of pipeline experiments that targeted a single set of cells over the course of approximately 2 weeks (5 different experiments with an average of 9.5 imaging sessions per experiment). Specifically, median adjustments of 16.47 μm (in X) and 16.34 μm (in Y) were required to match the 400 x 400 μm imaging FOV across sessions/days once the initial target FOV had been set on the first recording session. The right panel of Figure 7e shows these adjustments referenced to an example 2P FOV of GCaMP6+ neurons in the visual cortex, to further illustrate the relative size of these adjustments with respect to the fiducials (cells) within the FOV that are targeted across sessions.

Importantly, because the imaging plane is positioned normal to the putative orientation of the cranial window (and headframe ring), the X and Y image plane adjustments inherently consist of microscope stage movements in X, Y, and Z axes. Moreover, when considering the accuracy of this system it is important to note that the reported adjustments account for variability in tissue (cell) movement, stage movement, and headframe clamping variability— each of which are measured and defined in their own unique X, Y, Z coordinate spaces (see *System Alignment*). Thus, the image-plane X and Y adjustments, which are easily and quickly made by non-expert users, reflect the accuracy of the system to image a given set of cells in 3-dimensional space despite compounded sources of variability. In all, these systems exhibited variability well within our tolerances for performing longitudinal experiments targeted at a single set of GCaMP6+ cells within the mouse visual cortex.

## DISCUSSION

In order to build the *Allen Brain Observatory* pipelines that are capable of collecting standardized datasets from head-fixed mice over long-periods of time, we developed a series of integrated experimental platforms, each consisting of instruments that were built and registered to a shared coordinate space. Our cross-platform reference space strategy was based not only on creating a headframe and clamping systems, but also developing associated standard operating procedures for operation and routine monitoring and maintenance.

Creating a cross-platform reference space for our pipeline systems required three essential components: 1) a robust headframe, 2) a reproducible clamping system, and 3) data-collection systems that are built, and maintained, around precise alignment with a reference artifact. Additionally, the design of our pipeline systems had to meet the scientific requirements of our experimental goals and the teams of technicians who were responsible for operating the systems. Here we have described our head-fixation strategies for meeting the engineering, scientific, and operational requirements of our large-scale, *in vivo* pipelines.

We developed a headframe (and associated surgical tooling and clamping system) that met all of our pre-defined requirements pertaining to registration, rigidity, compatibility with our experimental systems, animal health, ease of handling, and future adaptability. The headframe design incorporated features that created three mutually intersecting perpendicular datum planes and, when used in conjunction with our clamping system, showed less than 4μm of deflection in our simulations, bench tests, and *in vivo* tests. We also observed that the titanium headframe is well tolerated by animals and allows for quick and easy handling and head-fixation by our technicians.

To integrate our head-fixation system into the *Allen Brain Observatory* pipeline we built a series of multi-modal data-collection systems that were all precisely aligned to a reference reticle that shares the same shank as the headframe. The precise alignment of these systems, and our ability to closely monitor alignment over time, allowed us to register our pipeline datasets across experimental platforms and leverage this registration for large-scale data collection. Here we report that these systems remain stable over long periods of time and frequent use and can be easily re-registered if deviations occur. Additionally, we describe an experimental application of cross-platform registration, made possible by our head-fixation system paired with routine system monitoring.

Lastly, our headframe is adaptable to other recording modalities and brain regions. Although our headframe was designed for our specific scientific goals, the mouse-interface portion (and associated surgical tooling) can easily be redesigned to accommodate alternative experimental modalities and/or recordings in other brain regions. Importantly, this can be done while still maintaining the mouse-to-screen geometry, which is accomplished by maintaining the design of the shank and its relationship to the skull fiducials. For example, we have adapted the initial Brain Observatory 2P headframe to gain access to more lateral visual areas as well as accommodate use of our multi-plane, Mesoscope imaging platform and objective (Supplemental Figures 6a and b, respectively). Additionally, we have adapted our headframe to accommodate different recording modalities including multi-probe electrophysiology, through-skull widefield imaging, and electroencephalogram (EEG) arrays (Supplemental Figures 6c-e). Although each of these headframe variants leverages the rigidity of the shank and clamping system, we have not yet performed deflection tests with these alternate versions and, as such, future adaptations of the headframe may require additional testing depending on experimental needs. Additionally, this headframe design is by no means compatible with all recording modalities and all brain regions. For example, the location of the headframe shank may not be compatible with recordings in the posterior region of the right hemisphere. As such, careful review of experimental requirements must be made when considering adopting components of this system.

The custom surgical tooling that we developed to standardize the surgical process was designed to interface with the off-the-shelf stereotaxic frames in our surgical facility. Specifically, the headframe clamp and levelling clamp interface with the KOPF Model #1900 dovetail block (on the stereotaxic arm) and earbar clamps, respectively. Therefore, additional modifications would be necessary to adapt these tools to other brands of stereotaxic frames (e.g., Narashige (https://usa.narishige-group.com). Importantly, however, because the conceptual approach of these tools remains the same regardless of brand, the only components that would require redesign would be those that directly interface with the stereotaxic frame (e.g., the dovetail block).

Our Allen Brain Observatory pipelines require head-fixation with access to the dorsal surface of the left hemisphere and therefore the systems we describe in this report are designed to be compatible with these conditions. In each of the experimental platforms, a technician interacts with the mice only to clamp the headframe onto the stage. It is worth noting that there have been a number of laboratories that have recently developed systems that allow mice to self-initiate head-restraint to allow 2-photon calcium imaging under stable conditions (e.g., Aoki, Tsubota, Goya, & Benucci, 2017; Murphy et al., 2016). Although these methods have the potential to provide high-throughput training while limiting handling by experimenters, currently these systems require two holding clamps and are therefore not compatible with our headframe design (see *Results*).

Since building the initial *Allen Brain Observatory* pipeline we have expanded the application of our engineering strategy to various other pipelines that utilize other recording modalities. Specifically, we extended the visual coding 2P pipeline to include a multi-depth 2P microscopy platform (Liu et al., 2019) as well as a multi-area/depth Neuropixels platform that is capable of recording from 6 high-density electrophysiology probes in an awake mouse (Siegle et al., 2019). In addition to our various “visual coding” pipelines, we have more recently built 3 “visual behavior” pipelines that allow for performing single- and multi-plane 2P (Tsyboulski et al., 2018), as well as Neuropixels, recordings from head-fixed mice performing visually-guided operant behavioral tasks (Groblewski et al., 2020). Each of the aforementioned *Allen Brain Observatory* pipelines were built around the cross-platform reference space described here and thus all meet our scientific and engineering requirements. In addition to the *Allen Brain Observatory* datasets being free to download from our web portal (https://portal.brain-map.org/explore/circuits), all files related to the tools and resources described in this report (including all headframe variants) are made freely available via our toolkit portal (https://portal.brain-map.org/explore/toolkit/hardware). Although it may not be feasible/desirable for external researchers to adopt our pipeline hardware and procedures *in toto*, we believe that incorporation of concepts and/or components of our engineering strategy could help to improve standardization and quality of physiology datasets obtained from head-fixed mice.

## MATERIALS & METHODS

### Headframe & Clamp Manufacturing

#### HEADFRAME

The headplate is manufactured from nominally 1.6 mm (.063 inch) thick titanium 6Al-4V annealed sheet and processed using common manufacturing methods. The Allen Institute selects material that is on the plus side of the sheet tolerance to prevent clamp adjustments that may be needed when switching between headplates that have thicknesses at the extremes of the tolerance band, which provides a maximum sheet thickness of 1.7mm (.068 inch). The raw sheet is waterjet cut, and the parts are tumble-deburred before being formed in a jig to add the correct angle for the mouse-interface. To prevent cracking during forming the component is heated slightly above room temperature to between 50 C and 100 C. Headplate manufacturing variances are kept to +/- 0.1 mm or better on all dimensions to ensure balance of manufacturability and compatibility with the clamping system, maintain consistency throughout the lifetime of the experiment, and to allow high precision placement of the headplate on the animal. It is worth noting that for smaller scale experimentation, headframes can be produced with 3D printing, without any compromises to datum surface registration or rigidity (Karolewska & Ligaj, 2019; Owsiñski & Niesłony, 2018). Additionally, if so desired, titanium headframes can be reused by removing them from mice post-mortem and cleaning with commonly used, titanium-compatible solvents such as acetone (Park et al., 2012). When produced in large quantities using subtractive processes the cost of each headframe is less than $15. Small quantities using additive processes results in a cost of approximately $125 per headframe.

#### CLAMP

The custom-designed clamping system components are all manufactured with common materials and traditional methods and can be produced on common CNC mills. The fasteners and alignment pins are commercially and commonly available. During assembly, alignment of the side clamp (Figure 5) is performed with custom tooling to ensure accuracy and repeatability of headplate clamping. All other components are pinned and aligned by reference features and no custom tooling is required. The clamp (including arm and base) can be produced for less than $500.

### Headframe Testing

#### SIMULATION

Design of the headframe and clamping systems were guided by assumptions on loading and the amount of mass a mouse could reasonably carry as an implant. With these basic assumptions the design effort proceeded with a goal to maximize stiffness of the headplate, while limiting the implant weight to 2 grams. Loads imparted into the system by a head-fixed behaving mouse were estimated to be at most .5N, or roughly an average weight mouse accelerating at 2 g vertically. External mechanical inputs were deemed to be negligible.

To compare designs and materials a static linear analysis was developed in Solidworks Simulation (Solidworks, Dassault Systemes). This simulation is founded on the “finite element method” and allows loads, supports and material behaviors to be realistically simulated when loaded in the linear elastic strain range of the materials involved, which is valid in this scenario. For this model an external vertically oriented load of 0.5N was distributed along the interior rim of the headplate (consistent with physical attachment to the mouse) and the clamp was constrained to zero motion at its fastener attachments. The headplate is modeled so as to be constrained to the clamp at the datum interfaces (Figure 2a). A solid mesh with approximately 1mm nominal element size was chosen to allow fine displacement detail to be resolved in the thin headplate and is well beyond the mesh density necessary to obtain results convergence.

The model was used to study and compare a variety of clamp and headplate materials including implantable plastics, aluminum, carbon fiber, stainless steels and titanium. The resulting design selected titanium 6Al-4V for the headplate and stainless steel 304 for the main clamping body. Results of this simulation are shown in Figure 3a and agrees well with bench testing of the loaded headplate as shown in Figure 3b.

#### BENCH TESTING

Bench deflection testing was performed using a Micro Epsilon laser triangulation displacement sensor, ILD1750-10 with a repeatability of 0.4 μm, a spot size of 110 μm and a range of 10mm. The instrument was placed within range, normal to the headplate at various testing locations. A static load was applied with calibrated weights freely hanging from the headplate from a small wire hook. The load was applied and released 3 times for a variety of different load scenarios and points of interest and is summarized in Figure 3b.

#### IN VIVO TESTING

Final deflection tests were performed on awake, locomoting mice that were head-fixed in our pipeline 2P microscope’s clamping system (Figure 3c). A Micro Epsilon sensor was used to first record baseline noise by recording deflection of a headframe alone (no mouse) clamped into the system (“Baseline”). Deflection was then assessed in 2 different mice that wereby obtaining multiple 20s recordings from the headframe surface (“MouseX_HF”) as well as from the surface of the cranial window (“MouseX_Win”).

### Surgery

All experiments and procedures were performed in accordance with protocols approved by the Allen Institute Animal Care and Use Committee. All surgeons received extensive training that combined hands-on training and a series of standard operating procedures pertaining to the use of both the custom and off-the-shelf surgical equipment (including skull leveling and skull fiducial identification). Headpost and cranial window surgery was performed on healthy male and female transgenic mice (p37-p63) weighing no less than 15 g at time of surgery and was based on a previously published protocol (Goldey et al., 2014). Pre-operative injections of dexamethasone (3.2 mg/kg, S.C.) were administered at 12h and 3h before surgery. Mice were initially anesthetized with 5% isoflurane (1-3 min) and placed in a stereotaxic frame (Model# 1900, KOPF; Tujunga, CA), and isoflurane levels were maintained at 1.5-2.5% for surgery. An incision was made to remove skin, and the exposed skull was levelled with respect to pitch (bregma-lambda level), roll and yaw. The stereotax was zeroed at lambda using a custom headframe holder equipped with stylus affixed to a clamp-plate (*see Headframe Surgical Tooling*). The stylus was then replaced with the headframe to center the headframe well at 2.8 mm lateral and 1.3 mm anterior to lambda. The headframe was affixed to the skull with white dental cement (C&B Metabond; Parkell; Edgewood, NY) and once dried, the mouse was placed in a custom clamp to position the skull at a rotated angle of 23° such that the visual cortex was horizontal to facilitate creation of the craniotomy (*see Headframe Surgical Tooling*). A circular piece of skull 5 mm in diameter was removed, and a durotomy was performed. A glass coverslip (cut from a single piece of glass to obtain a “stacked” appearance that consisted of a 5 mm diameter “core” and 7 mm diameter “flange”), was cemented in place with Vetbond (3M; St. Paul, MN). Cement was then applied around the cranial window inside the well to secure the glass window. Post-surgical brain health was documented using a custom photo-documentation system and animals were assessed one, two, and seven days following surgery for overall health (bright, alert and responsive), cranial window clarity and brain health.

## ACKNOWLEDGEMENTS

We thank Michael A. Buice, Marina Garrett, Leonard Kuan, Shawn Olsen, Carol Thompson, Wayne Wakeman, Jack Waters, Derric Williams, Nathalie Gaudreault, Fiona Griffin, Perry Hargrave, Robert Howard, Josh Larkin, Rachael Larsen, Eric Lee, Arielle Leon, Jennifer Luviano, Thuyanh Nguyen, Jed Perkins, Miranda Robertson, Sam Seid, Casey White, Ali Williford, Hongkui Zeng, Amy Bernard, John W. Phillips, R. Clay Reid, and Christof Koch for their intellectual and/or experimental contributions.

We thank the Animal Care, Transgenic Colony Management, Lab Animal Services and Neurosurgery & Behavior teams for mouse husbandry, care, and surgery.

We thank Allan Jones for providing the critical environment that enabled our large scale team effort. We thank the Allen Institute founder, Paul G. Allen, for his vision, encouragement, and support.

---

**Supplemental Figure 1.**
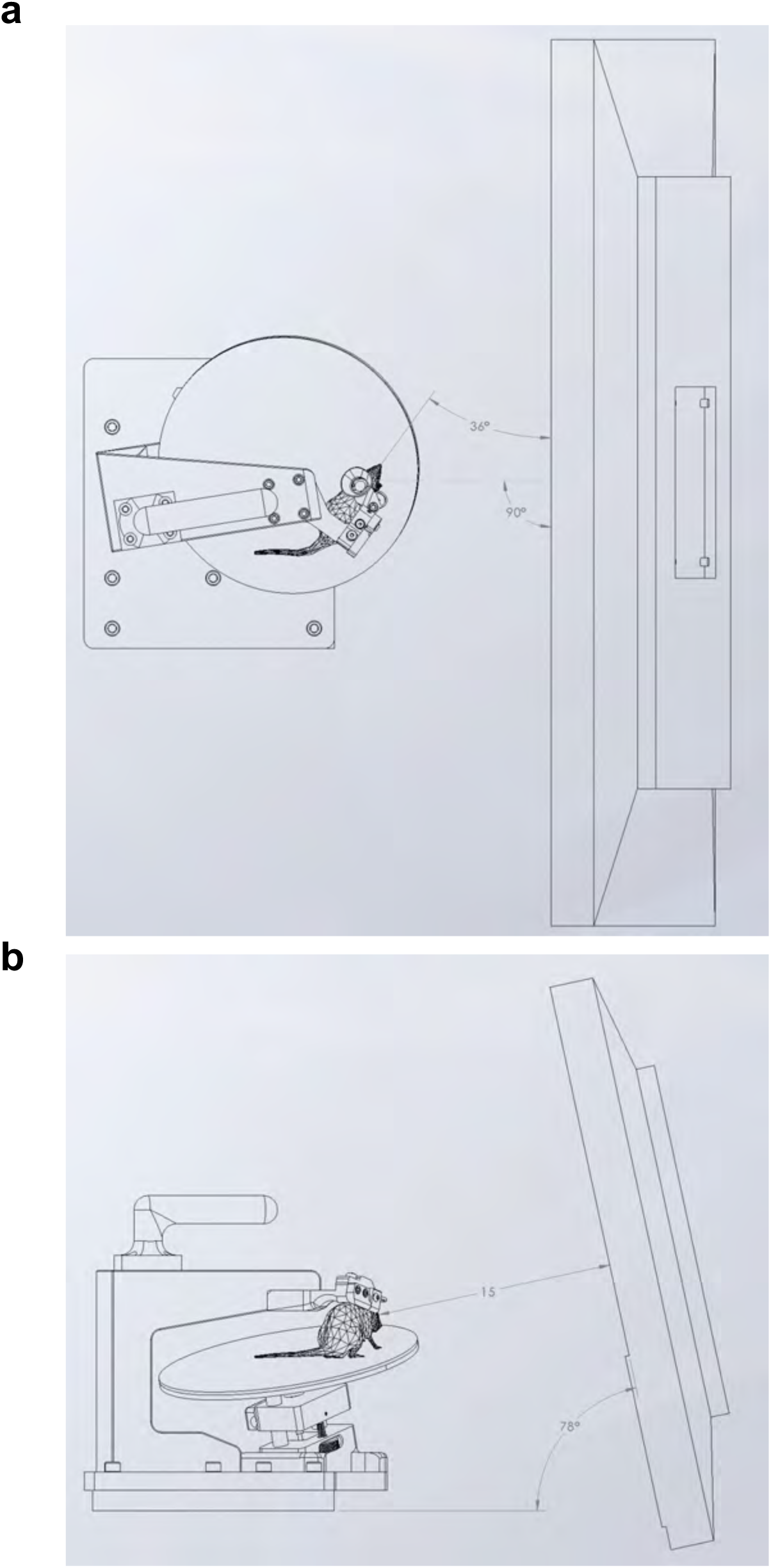
Brain Observatory pipeline mouse-to-screen geometry. a) Top-down view of the pipeline mouse-to-screen geometry. Mice are positioned at a 36° angle with respect to the screen. b) Rear-view of the pipeline mouse-to-screen geometry. Mice are positioned 15cm from an LCD monitor, which is positioned at a 78° angle with respect to ground.

**Supplemental Figure 2.**
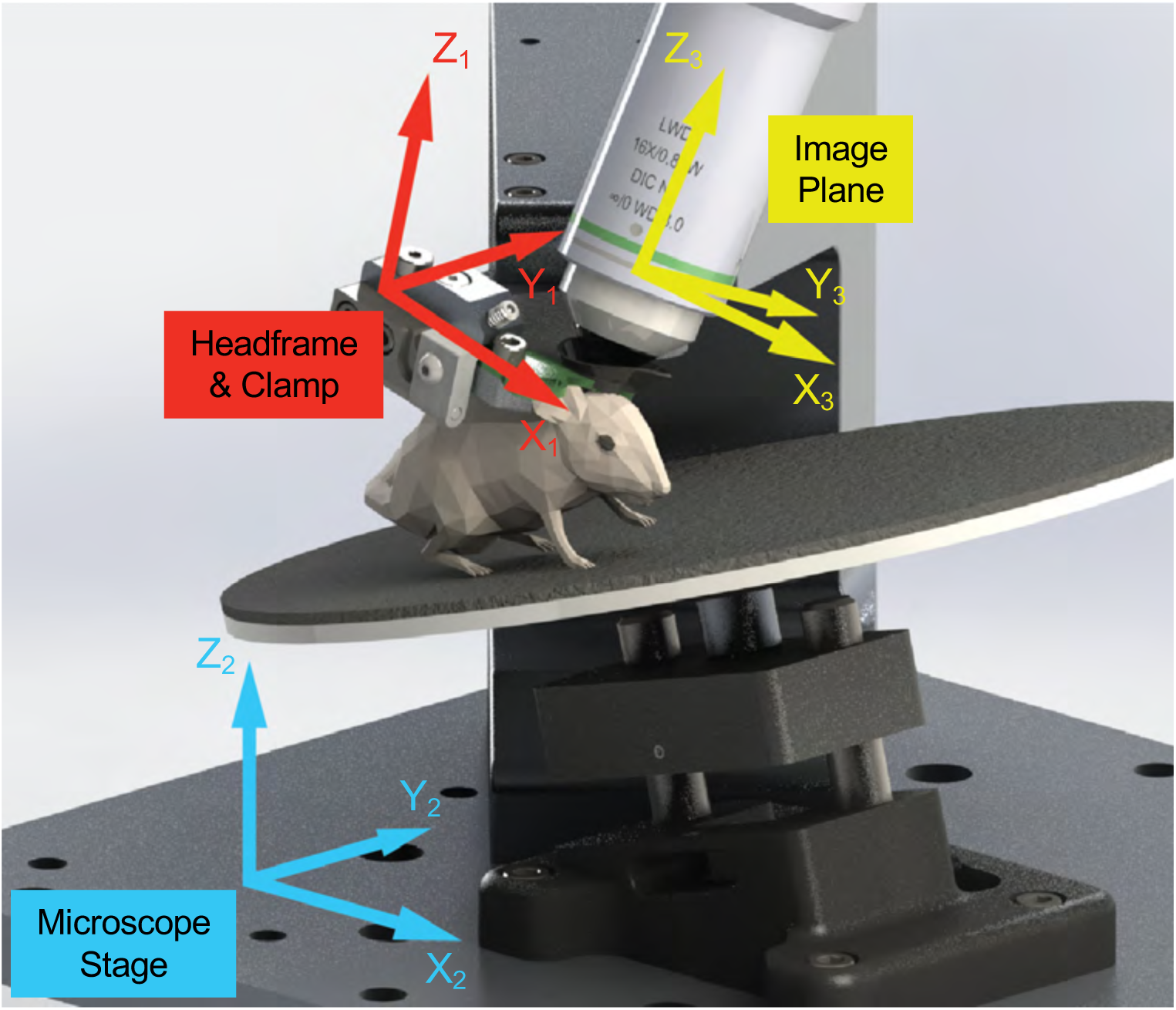
Representation of system coordinate geometries. Each pipeline system (such as the 2P microscope platform shown here) consisted of multiple coordinate spaces. As such, 3-dimensional movements and/or variability in one coordinate space (such as the headframe clamping system) resulted in a different, 3-dimensional shift in the other coordinate spaces. In order to facilitate FOV targeting in imaging space, systems were built, aligned, and maintained using a reticle system.

**Supplemental Figure 3.**
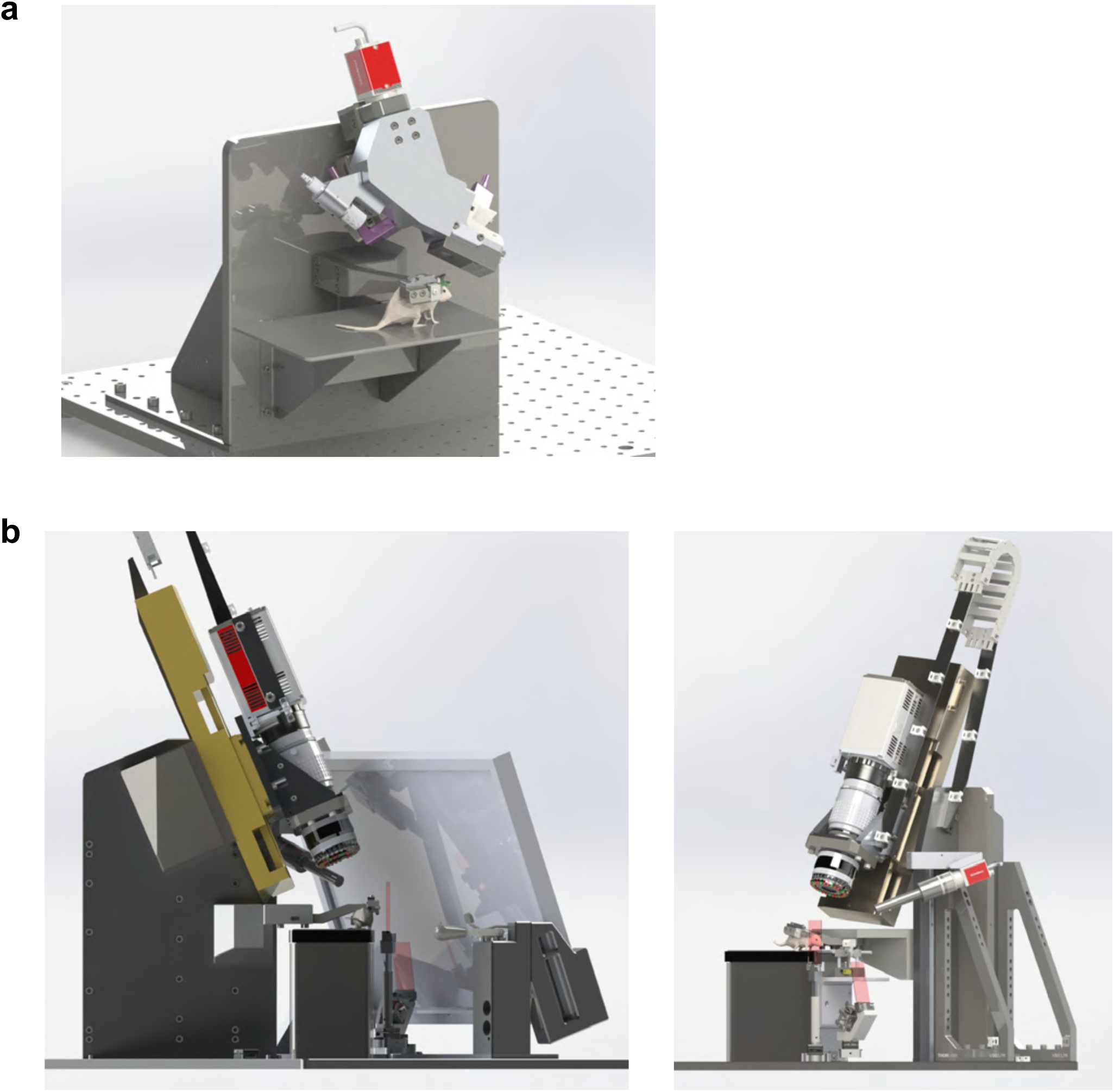
Surgical photo-documentation and Intrinsic-Signal Imaging pipeline systems. a) The surgical photo-documentation system was used to acquire registered images of the cranial window immediately following surgery. No visual stimulation was required. b) The intrinsic-signal imaging systems (shown with and without the screen placed at a modified mouse-to-screen geometry) were used to acquire individualized maps of functional boundaries between visually-responsive regions of the mouse cortex. Systems included a camera (with ring-light LED illumination), eye-tracking camera (with LED illumination), and anesthesia machine (Somnosuite)

**Supplemental Figure 4.**
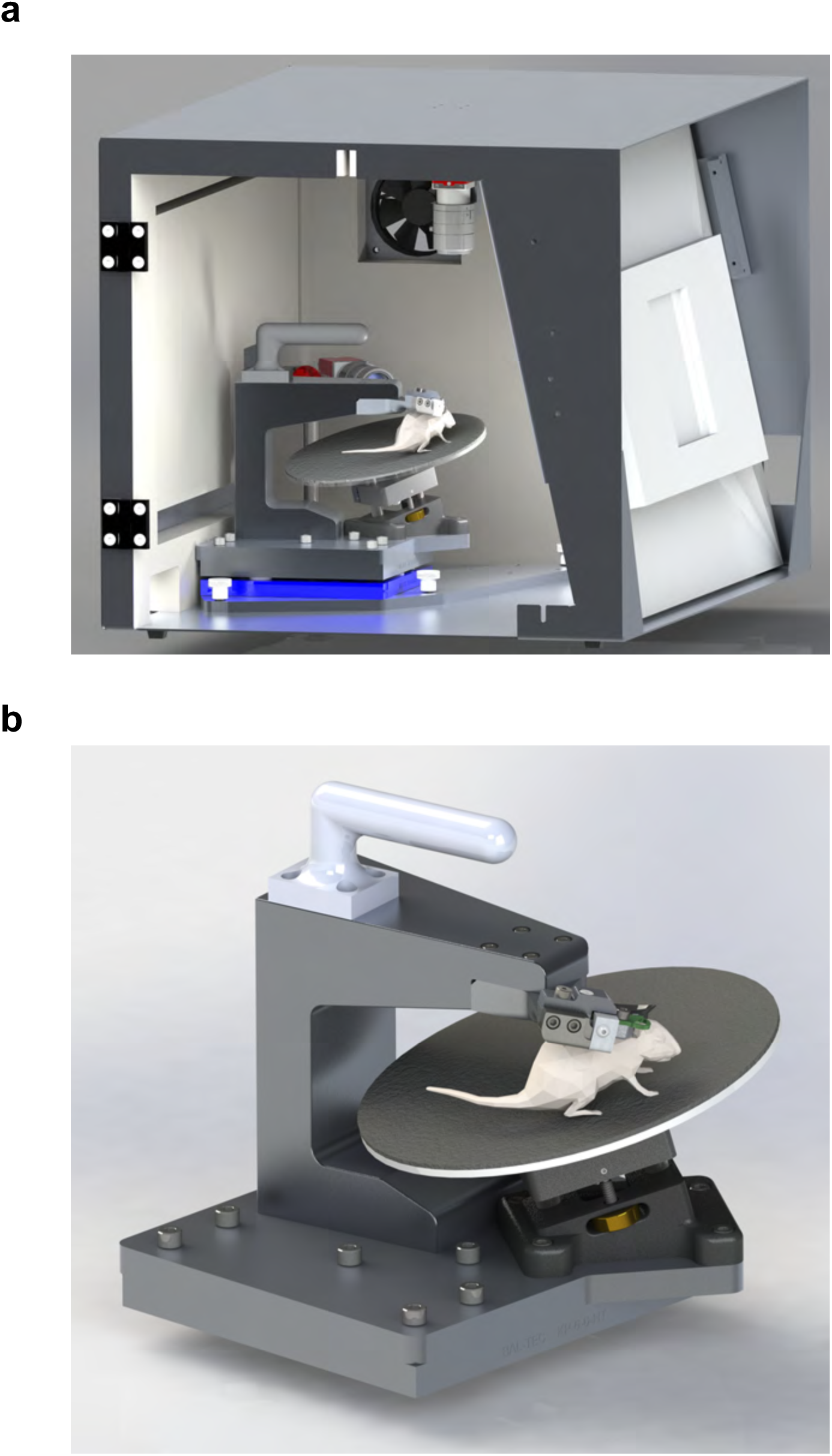
Behavioral enclosure and removable mouse head-fixation stage. a) Behavioral enclosures allowed for mice to be habituated to head-fixation and trained on visually-guided operant tasks. Front view of an enclosure (without door) shows a removable stage precisely positioned with respect to the stimulus screen using a kinematic mount (in blue). Enclosures are equipped with sound-attenuating foam, ventilation fan, body camera with illumination source, and a fluid delivery system mounted to a motorized 3-axis stage (not pictured). b) The removable mouse stage allowed for mice to be head-fixed outside of the enclosure, an important factor in making the enclosures compact in size.

**Supplemental Figure 5.**
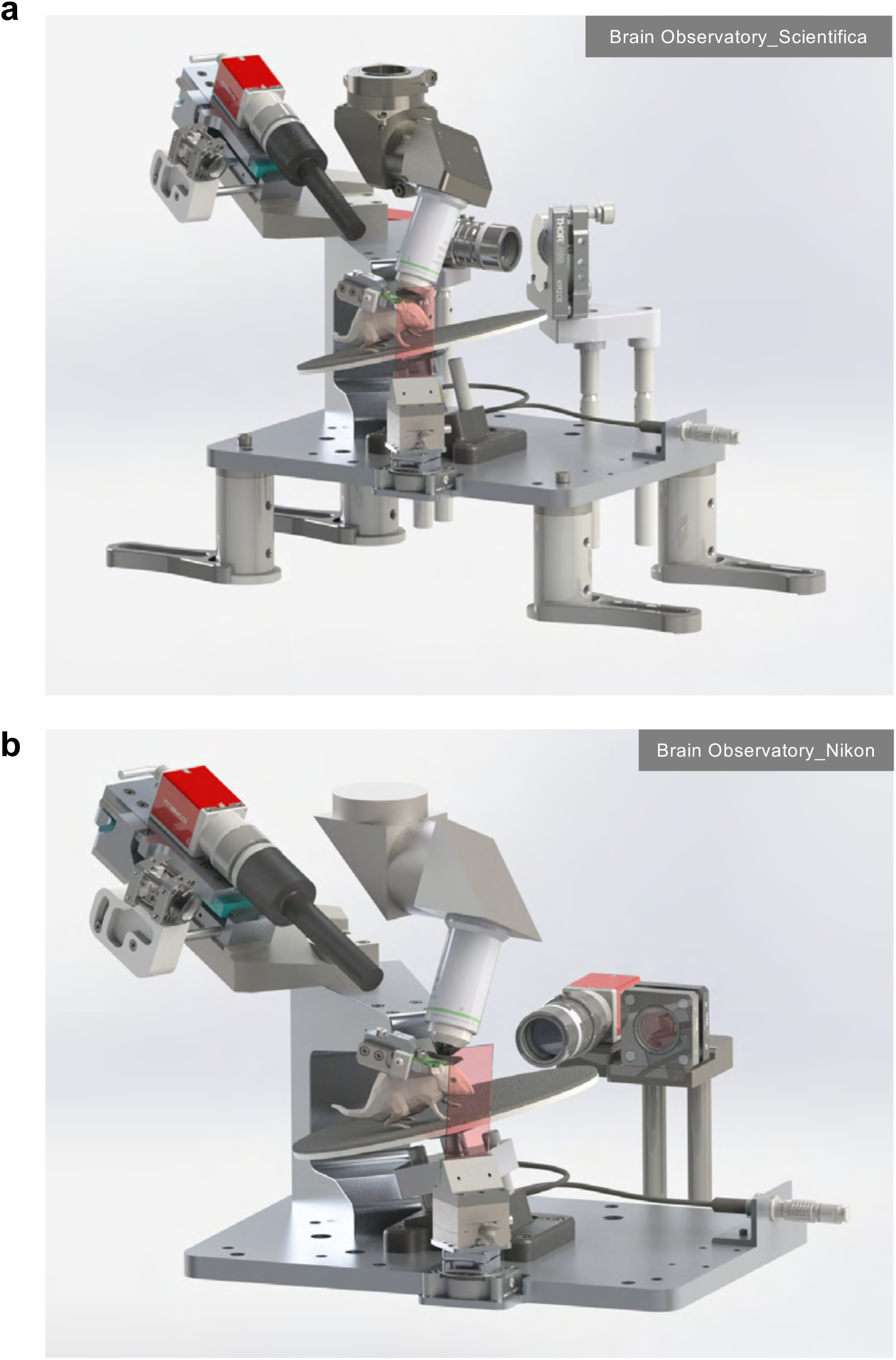
Pipeline 2-photon microscopy platforms. a) Scientifica Vivoscope microscope (equipped with a 16X Nikon CFI LWD Plan Fluorite objective) with custom modifications to accommodate Brain Observatory mouse-to-screen geometry. b) Nikon A1R MP+ microscope (equipped with a 16X Nikon CFI LWD Plan Fluorite objective) with custom modifications to accommodate pipeline mouse-to-screen geometry.

**Supplemental Figure 6.**
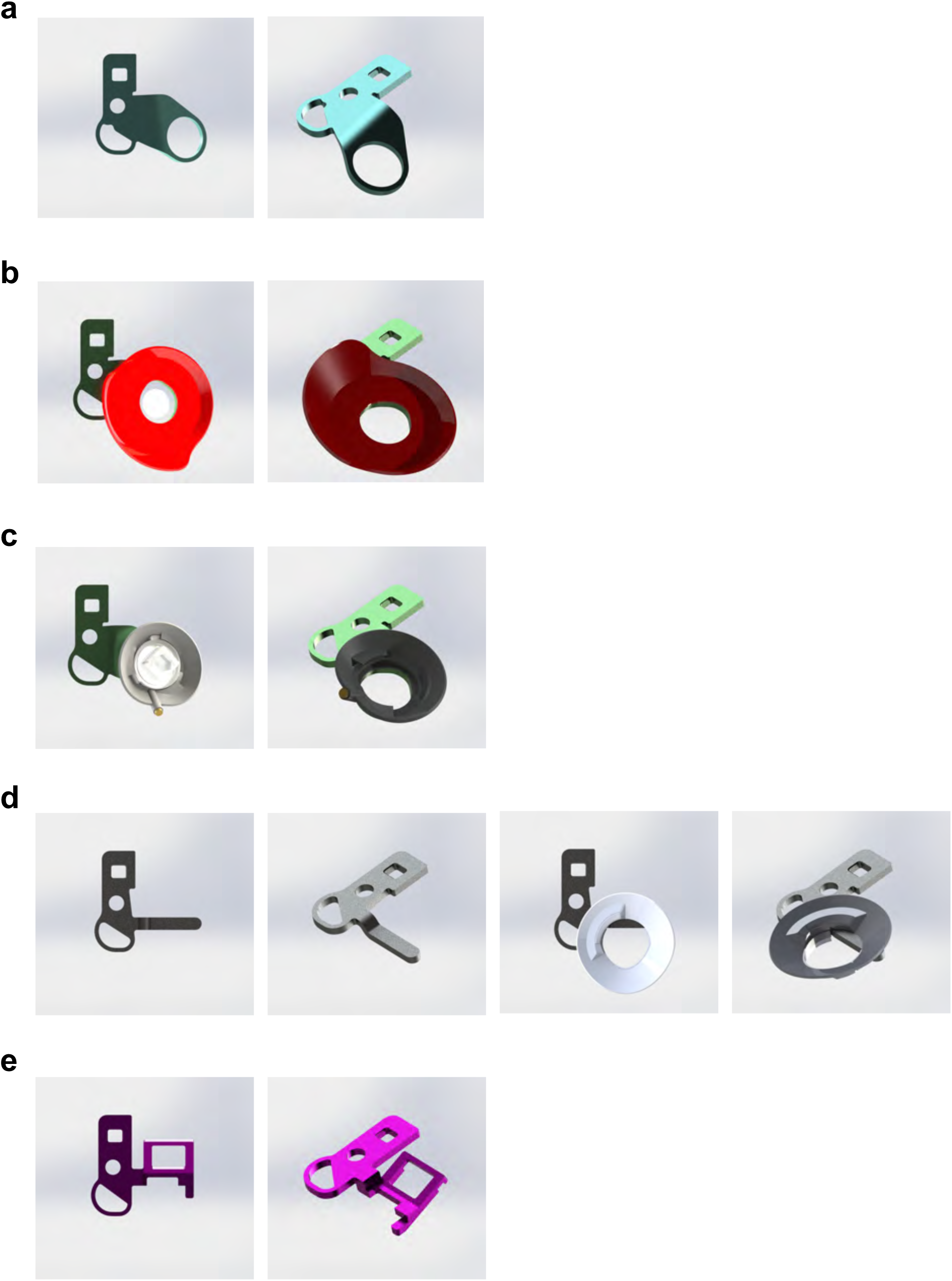
Brain Observatory headframe variants. a) Headframe used for 2P imaging from lateral visual areas. b) Headrame and well used for 2P imaging with the Mesoscope microscope and objective. c) Headframe, well, and cap used for multi-probe electrophysiology recordings with Neuropixels probes. d) Headframe and well used for either through-skull, widefield imaging or stereotaxically-guided probe insertion. e) Headframe used for electroencephalogram (EEG) arrays.

